# Optically driven control of mechanochemistry and fusion dynamics of biomolecular condensates via thymine dimerization

**DOI:** 10.1101/2025.03.30.646143

**Authors:** Vahid Sheikhhassani, Faith F.H.K. Wong, Daniel Bonn, Jeremy D. Schmit, Alireza Mashaghi

## Abstract

Phase-separated biomolecular condensates serve as functional elements of biological cells, contribute to protocell formation in prebiotic systems during early life, and represent a distinct class of soft matter with a broad range of potential applications. Understanding and controlling condensate mechanochemistry is critical for their function and material properties. Photochemical processes, such as UV-induced chemical modifications, are ubiquitous in nature and can have both detrimental and constructive impacts on living systems, and are also readily implemented in engineering applications. However, how phase-separated condensate formation influences photochemical processes, and conversely, how photochemical reactions impact condensate dynamics, remains an open question. Combining scanning probe microscopy with optical imaging and control, we developed assays that enable the study of mechanical transitions and fusion dynamics in condensate droplets, revealing that UV-induced thymine dimerization alters condensate nucleation and coalescence. Depending on the frequency and topological arrangement of thymine dimers, particularly the balance between inter- and intrachain crosslinks, UV can induce a transition from liquid-like to solid-like behaviours or lead to aggregate formation. UV treatment also leads to compartmentalization in condensate systems by e.g., promoting the formation of arrested fusion droplets, which are stable against environmental changes. UV illumination can thus be leveraged to program the architecture and material properties of DNA-based biomolecular condensates, with implications for prebiotic chemistry, and bio-inspired engineering.

## Introduction

Biomolecular condensates formed through liquid-liquid phase separation (LLPS) play key roles in various cellular processes, ranging from gene regulation to stress granule formation ^1–5^. These condensates assemble through multivalent interactions among macromolecules such as nucleic acids and proteins^6–9^, and while physiological condensates are often liquid-like, they can exhibit diverse material and mechanical properties, particularly upon aging or under stress conditions that alter their interactions^10–15^. The strength, density, and topological arrangement of interactions, along with the intrinsic mechanical properties of the polymer chains, are critical determinants of condensation and the resulting mechanical phenotype^16^. Such variations in condensate material state may directly influence molecular transport and chemical reactions within their microenvironments, effects central to cellular biochemistry and exploitable for reaction control in engineered systems^6,17–21^. Modulation of the mechanical properties of condensates is typically achieved by introducing new molecules (e.g., crosslinkers) or adjusting molecular concentrations, approaches that inherently alter condensate composition, making it difficult to disentangle the effects of mechanics and chemistry^22^.

Light can be exploited to modulate material properties in soft and polymeric systems^23,24^. Yet, light-triggered control over the mechanical state of condensates, achieved without significantly altering their chemical composition, offers a largely unexplored route for engineering condensates. One particularly relevant form of light stimulus is ultraviolet (UV) radiation, which is both a major cause of DNA damage through the formation of covalently bonded pyrimidine dimers such as thymine dimers (TT) and a critical factor in the emergence of life on Earth^25,26^. TT dimers disrupt normal DNA replication^27^ and transcription^28^, leading to mutations that can initiate various diseases, and are the primary cause of skin cancer^29^. Cells recognize and attempt to repair these lesions^30^; however, persistent or improperly repaired TT dimers can give rise to short stretches of single-stranded DNA (ssDNA), ultimately contributing to genomic instability and carcinogenesis^31^.Furthermore, UV radiation was abundant on early Earth and contributed to the synthesis of building blocks of life^32^, while coacervate condensates of these building blocks are also considered plausible protocellular structures^33^. The droplets feature a highly dynamic molecular environment^34^, and are permeable to molecules from their surroundings with certain degrees of selectivity, enabling chemical synthesis and molecular recognition (e.g., receptor-ligand interactions) required for life^35^. Nucleic acids, known for their propensity to form condensates, were likely key components of protocells^33^. Because coacervates lack lipid membranes, these protocellular structures provide only limited shielding of their contents from solar UV. It remains unclear, however, what UV does to (nucleic acid) coacervates and whether it can shape their structural mechanics.

A technical challenge in studying the mechanochemistry of condensate droplets is the difficulty in probing viscoelastic and fusion properties as they undergo liquid-to-solid transitions. Active microrheological characterization of condensate droplets is mostly performed utilizing optical tweezers (OT)^36,37^, which are constrained to low-force regimes and generally rely on trapped beads. Micropipette aspiration is also constrained by achievable force (suction pressure) and experimental throughput^38,39^. Recently, scanning probe microscopy (SPM), a technique widely used in mechanobiology and materials science^40^, has been applied to characterize the material properties of condensates^22,41,42^, covering a significantly higher range of forces and eliminating the need for exogenous particles in the system. Yet, SPM has never been applied to study droplet-droplet fusion, a gap that, if addressed, could enable unprecedented insights into condensate mechanics and interactions.

Here, we introduce a photochemical strategy to precisely modulate condensate mechanics using UV-induced thymine dimerization, enabling control over mechanical properties while preserving chemical composition —that is, without adding new molecules and without changing atomic composition. Using poly-thymine single-stranded DNA (dT40) and poly-L-lysine (PLL) as a model system, we demonstrate how this strategy drives transitions among distinct mechanical states, from liquid-like droplets to more rigid, gel- or solid-like networks. To capture these mechanical changes, we developed SPM-based assays combined with optical imaging for label-free characterization of condensate mechanics and fusion dynamics. These transitions lead to dramatic changes in fusion behaviour, ultimately driving compartmentalization, a process relevant to prebiotic chemistry and bio-inspired materials design. We attribute these observations to the formation and topological arrangement of UV-induced TT dimers, which can be local, long-range, or occur between chains. Our study provides a compelling example in which a photochemical reaction alters the condensed phase, and conversely, condensation modulates photochemical reaction outcomes.

## Materials and Methods

### Preparation of ssDNA and Poly-L-Lysine Stock Solutions

ssDNA oligo of dT40 was purchased from Integrated DNA Technologies (NJ, USA). The dry stocks were reconstituted in RNase-free water with no added salt. DNA concentration was measured subsequently using a Qubit spectrophotometer. The DNA stocks were then aliquoted and stored at −20°C until use. Poly-L-lysine (PLL) hydrochloride with an average molecular weight of 15,000 to 30,000 Da was used in all experiments. The PLL was dissolved in water to prepare stock solutions at 22.4 mg/ml. Based on the average molecular weight of the PLL, the average length of the polymer is estimated to be 150 residues long. This range of lengths serves as a simple model for the diversity of positively charged polypeptides found in cells or in primitive life forms.

### Surface passivation for rheological and droplet-droplet interaction analysis

Condensate samples for phase contrast imaging, mechanical analysis, and droplet-droplet interaction analysis were prepared on TPP Petri dishes. For rheological analysis, dishes were passivated with 1% w/v BSA (A2153) solution, incubated for 30 min, and washed five to six times using RNase-free water. For droplet-droplet interaction analysis, the experimental setup required detaching a droplet from the substrate using the cantilever and positioning it on top of a target droplet for the fusion assay. A combination of Pluronic and BSA passivation was used to enable this experiment. On the Pluronic-coated surface, droplets were loosely bound, allowing easy detachment, whereas on the BSA-coated side, droplets showed stronger adhesion, preventing detachment during the fusion process and enabling complete coalescence on the surface. A guideline dividing the dish in half was drawn on the underside of the Petri dish to ensure consistent application of the passivation agents. The dish was securely fixed in position and tilted to approximately 20 degrees. For each passivation, the bottom half of the dish was filled with the corresponding passivation agent up to the guideline, incubated for 30 minutes, and washed multiple times by carefully adding RNase-free water.

### Sample preparation

Each time, for sample preparation, 60 μL of the working buffer (150 mM KCl (p9333), 10 mM imidazole (56750), and 0.01% w/v NaN_3_ (8.22335))^43^ was placed as a round droplet at the center of the dish. For droplet-droplet analysis, this step was performed carefully to align the droplet’s middle axis with the drawn guideline. To mimic the intracellular environment, the crowding agent Ficoll® PM 70 (F2878) was added to the buffer at a final concentration of 50 g/L. Subsequently, ssDNA and poly-L-lysine (PLL, P2658) were added sequentially to reach a final concentration of 30 µM for each. After adding each component, the solution was gently mixed by pipetting 10 times.

### SPM measurements

A JPK CellHesion 200 (Bruker, Germany) equipped with a phase contrast microscope delivering real-time *in-situ* images of measurements utilizing a 20X/0.4 objective was employed in this study. Mechanical analysis was performed as explained by Naghilou et al.^22^ Briefly, a 60 µl bulk drop containing the condensates was prepared, and the droplets were allowed to settle on the dish. All experiments were performed within 4 h of droplet formation. Before each mechanical measurement, a 5 µl drop was carefully taken from the top of the bulk droplet and placed on the cantilever to avoid the generation of air bubbles when the head is placed. As it has been shown that micrometre-scale indenters lead to a better signal-to-noise ratio in comparison to their nano-scale counterparts^44^. For all experiments, SAA-SPH-5UM cantilevers (Bruker, Germany) made of Si_3_N_4_ with a hemispheric tip (23 µm height, 5.13 µm radius) were used. The calibration of the amplitude and the exact cantilever spring constant was determined with the thermal noise method^45^. The cantilever was passivated with 1% Pluronic F127 (P2443) for 30 min to prevent the adhesion of droplets on the tip after measurements^46^. Condensates (preferably of 8-10 µm in diameter) were indented with a 0.3 nN set force and a cantilever approaching with a constant velocity of 1 µm/s. Acquired data was post-processed by Bruker data analysis software and prepared for calculation of rheological parameters using a custom Mathematica code (Mathematica 14.0, Wolfram). Size distribution analysis and calculation of the radius of condensates were done using ImageJ (https://imagej.nih.gov/) from the phase contrast images.

### Confocal fluorescence imaging

Confocal fluorescence imaging was performed using a Nikon Ti2 inverted microscope with a Nikon C2plus confocal system and a Plan Apo VC 20x DIC N2 objective (NA = 0.75, WD = 1000 μm). Imaging was conducted with a Galvano scanner in one-way scan mode at 6.897x zoom. Excitation was achieved with 489 nm and 562 nm lasers for GFP and TYE-665, respectively, with emissions detected at 540 nm and 665 nm. The detector gain was set to 100 (GFP) and 70 (TYE-665), and the pinhole size was 40 μm. A brightfield transmission detection (TD) channel was also acquired. Images (512 × 512 pixels) were acquired in three planes: GFP, TYE-665, and TD, with an exposure time of 1095.456 ms at a voxel size of 1×1×1 pixel³. Image processing and analysis were performed using Fiji (ImageJ) version 2.14.0/1. TIFF-formatted images were used for quantitative and structural analysis without additional thresholding or scaling.

### UV–visible spectroscopy

UV–visible spectroscopy was performed to evaluate photochemical modifications to the dT40 oligonucleotide following UV irradiation. A final concentration of 30µM dT40 was used for all experiments. For the +UV45_Premix samples, the DNA was mixed into 60µl of working buffer and irradiated for 45 minutes, after which a sample was taken for spectroscopic analysis. For the +UV45_4h samples, 45 minutes of irradiation were applied after 3 hours and 15 minutes of the condensate formation period. Subsequently, high salt concentration was added to the droplets to release the dT40 into the dilute phase, from which a sample was collected. Non-irradiated control samples were prepared by mixing dT40 in the working buffer and analysed immediately. Absorption spectra were recorded using an Agilent 8453 UV–Vis spectrophotometer (Agilent Technologies, Santa Clara, CA, USA) in a low-volume quartz cuvette across the standard nucleic-acid absorption range. The resulting spectral changes were used to identify UV-induced TT dimerization in dT40.

### Microfluidic imaging (MFI)

Particle imaging was performed using a ProteinSimple MFI system (ProteinSimple, San Jose, CA, USA) operated in basic mode using default acquisition parameters with a 200 µl/minute flow rate. Particle identification and quantification, including particle counts, equivalent circular diameter (ECD), and circularity, were carried out with the instrument’s analysis software (MFI View System Software, ProteinSimple, San Jose, CA, USA). Processed datasets were exported and plotted using OriginLab 2025 (OriginLab Corporation, Northampton, MA, USA). The approach was applied to +UV45_0h samples, right after UV illumination, and the size and morphology of the floating objects were quantified.

### Fluorescence recovery after photobleaching (FRAP)

FRAP experiments were performed to quantify the relative recovery dynamics of PLL and DNA within individual condensate droplets. To enable simultaneous measurement of both components, 5% FITC-labeled PLL (P3543; Sigma-Aldrich, St. Louis, MO, USA) was mixed with its unlabeled counterpart, and 5% TYE-665–labeled dT40 was incorporated into the DNA component. Sample preparation followed the procedures described above.

All measurements were carried out on a Nikon Eclipse Ti2 microscope (Nikon Instruments, Tokyo, Japan) equipped with a 20×/0.75 NA objective. A circular region of interest (radius 1.25 μm) within each droplet was bleached for 1 s using a 488 nm laser at 250 μW, ensuring that droplet diameters were at least two to three times larger than the bleach radius. Fluorescence recovery was recorded for 3 min at low excitation power to minimize additional photobleaching. Both fluorescence channels (FITC for PLL and TYE-665 for DNA) were acquired sequentially for each droplet. Fluorescence intensities were extracted from the bleached region (*F*_ROI_), an unbleached reference droplet (*F*_ref_), and a background region (*F*_bkgd_). To correct for photodecay, intensity traces were normalized to the reference droplet according to:

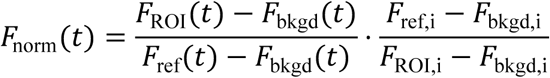

where *F*_i_ denotes the pre-bleach average intensity. Normalized recovery curves were used to compare relative recovery times of PLL and DNA within the same droplets.

### Microscale thermophoresis

Microscale thermophoresis (MST) was used to assess the mobility of dT40 molecules in PT40, +UV45_Premix, and +UV45 samples. All samples were prepared at a final dT40 concentration of 30 µM. For the +UV45_Premix condition, dT40 was added to 60 µL of the bulk solution, irradiated for 45 min, and samples were collected immediately after irradiation. For the +UV45 condition, droplets were allowed to form for 3 h 15 min before 45 min of UV exposure. High-salt buffer was then added to dissolve the condensates and release dT40 into the dilute phase, from which material was collected. Non-irradiated controls were prepared by mixing dT40 directly into the working buffer and measuring immediately.

MST measurements were performed on a Monolith NT.115 instrument (NanoTemper Technologies, Munich, Germany) using standard capillaries. Samples were equilibrated for 10 min at room temperature before loading. Fluorescently labeled DNA (5% TYE-665 dT40) was mixed with unlabeled samples. Thermophoretic traces were recorded at 25 °C using medium IR-laser power. Normalized fluorescence (F_norm_) was extracted using the manufacturer’s software, and changes in F_norm_ were used to quantify differences in ssDNA mobility associated with UV-induced TT dimer formation.

### Anti-CPD immunofluorescence staining assay

Cyclobutane pyrimidine dimer (CPD) formation within condensates was assessed by immunofluorescence staining using a FITC-conjugated anti-CPD antibody from the OxiSelect™ Cellular UV-Induced DNA Damage Staining Kit (CPD, Cell Biolabs). Condensate samples contained 5% TYE-665–labeled DNA (of the total DNA) and were incubated with the anti-CPD primary antibody diluted 1:100 in assay diluent for 1 h at room temperature with gentle agitation. After three washes with 1× wash buffer, droplets were incubated with a FITC-conjugated secondary antibody (1:100 in assay diluent) for 1 h under identical conditions. Excess antibody was removed by four additional washes with 1× wash buffer. Fluorescence microscopy was performed using channels appropriate for both TYE-665 and FITC fluorescence detection.

## Results

### UV irradiation affects the phase separation behaviour and promotes arrested fusion droplets

We first explored the phase diagram of PLL/dT40 by varying component concentrations and selected a 1:1 PLL:dT40 ratio at 30 µM of each component for subsequent analyses (Figures 1A and S1), as this condition produced micron-sized droplets that were well suited for mechanical measurements and imaging. Prior to investigating the effects of UV irradiation, we characterized the baseline phase separation behaviour of PLL/dT40 under varying salt conditions, from 0 M to 1 M. No droplets formed at either extreme: high salt screens electrostatic interactions below the threshold for phase separation, while low salt favors tight 1:1 heterodimers with a strong positive net charge (due to charge asymmetry −40 vs. +150), preventing further association. Intermediate salt concentrations, by contrast, allow molecules to exchange partners within a liquid environment^47,48^, promoting spherical condensates of varying size. The largest droplets were consistently observed at 150 mM KCl (Figure 1B and C), a condition used for the remainder of our study.

**Figure 1.**
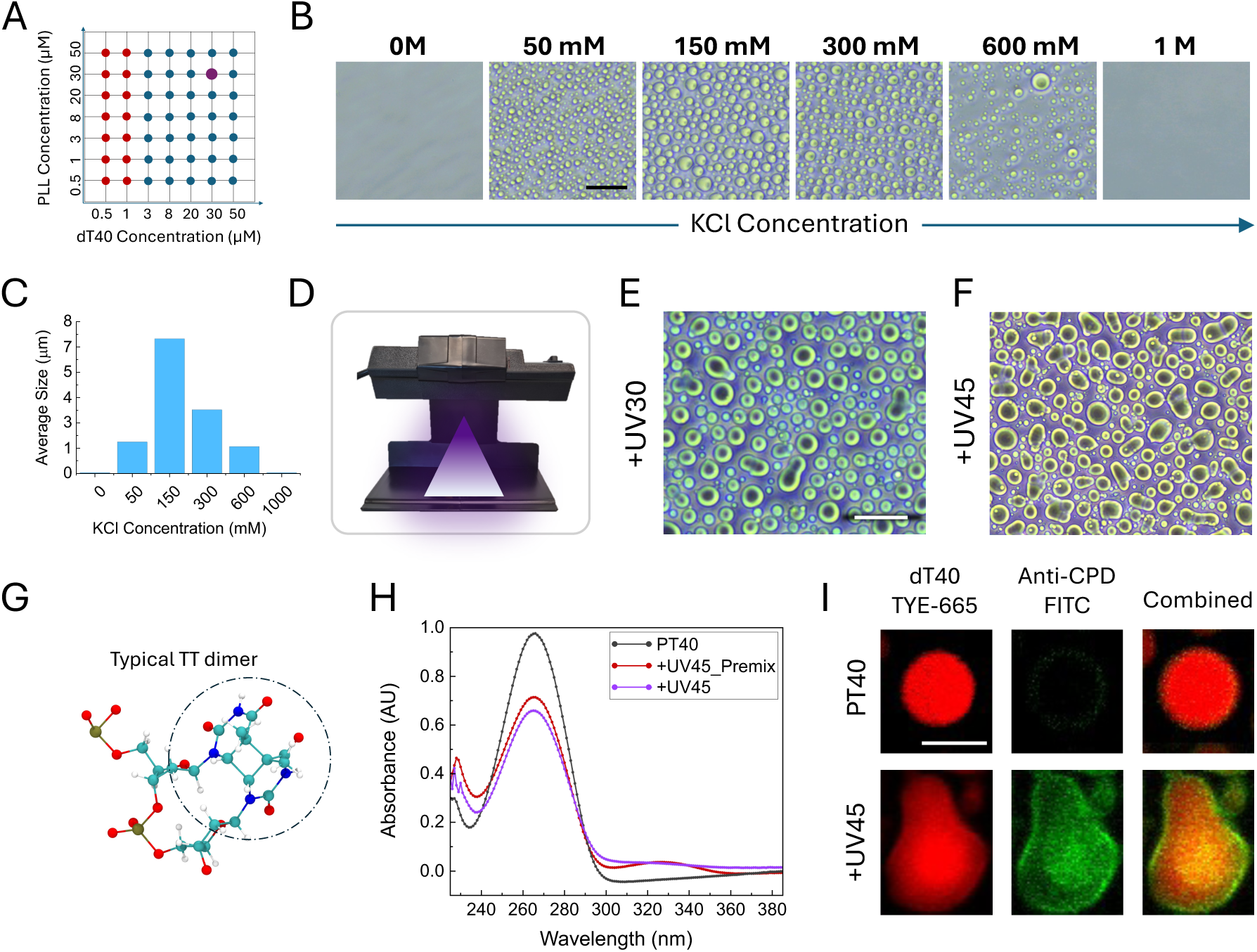
Morphological and chemical characterization of PLL:dT40 condensates with and without UV illumination. A) Phase diagram illustrating condensate formation at different PLL and dT40 ratios. Blue dots represent conditions where droplet formation was observed, while red dots indicate no droplet formation. A selected condition (30 µM PLL: 30 µM dT40) highlighted in purple was used for all the preparations. B) Effect of salt concentration on condensate formation. No condensates formed at extreme salt concentrations (0 M and 1 M KCl), while intermediate KCl concentrations resulted in the formation of droplets. C) Quantification of average droplet diameters at different salt concentrations. The highest average droplet diameter was recorded at 150 mM KCl. D) UV irradiation setup. UVC lamp installed on a modified stand ensured uniform exposure. All samples were placed in a fixed position on the holder beneath the lamp. Images were captured after a total of 4 hours of sample preparation, including 30 and 45 minutes of UV irradiation at the final stages. E,F) Phase-contrast microscopy showed clear morphological changes in the droplets after 30 and 45 minutes of UV exposure, with elongated droplets appearing after UV exposure. These elongations were more pronounced following 45 minutes of irradiation. G) Ball-and-stick representation of a typical TT dimer(depicted from PDB structure 9j8w). H)UV-vis absorption spectra of dT40 before and after UV exposure. The non-irradiated dT40 (black line) displays the characteristic absorption maximum near 260 nm, corresponding to the π-to-π* transitions of the thymine bases in ssDNA. Following UV exposure, both spectra exhibit a hypochromic effect, a pronounced decrease in absorbance at 260 nm, indicating the formation of thymine-thymine (TT) cyclobutane dimers and the associated disruption of the electronic environment of the bases. Furthermore, a weak shoulder appears between 300 nm and 320 nm in the irradiated spectra, consistent with the formation of minor photoproducts, such as the (6-4) photoproducts or their Dewar isomers. Critically, the reduction in 260 nm absorbance and the intensity of the 300-320 nm shoulder are more pronounced in the +UV45 (after condensate formation) sample compared to the +UV45_Premix sample (before mixing with PLL). I) Immunofluorescence reveals the formation of TT dimers within UV-irradiated condensate droplets. To directly probe the chemical change, condensates were stained with an anti-TT dimer (Anti-CPD) antibody (FITC-labelled). The dT40 samples (top row) confirm ssDNA assembly (red) but show no CPD signal (green). In contrast, the +UV45 sample (bottom row) displays a strong CPD signal co-localized with the dT40 (yellow/orange combined image), providing direct chemical proof of TT-dimer formation within the bulk of the condensate. The scale bar represents 20 µm for phase-contrast images and 5 µm for confocal images.

For UV treatment, samples were prepared over a total period of 4 hours and placed under a UVC lamp (UVP UVG-11, λ=254 nm; 1120µW/cm² at the sample) (Figure 1D), with UV illumination applied during the final 30 (+UV30) or 45 (+UV45) minutes^49^. Spectroscopic analysis indicates that UV induces TT dimerization in dT40 under this exposure condition (Figure 1G, and H). Although the UV illumination times used in this study are much longer than the picosecond timescale required for the formation of intramolecular TT dimers^50^, it is comparable to the timescale characterizing intermolecular crosslinking^49^. Phase-contrast microscopy immediately after irradiation revealed altered droplet morphologies with apparent reduction in circularity (Figure 1E, and F). Similar UV-induced morphological changes were observed with droplets formed on nonsticky surfaces (Figure S2), suggesting that observed patterns are primarily driven by UV-induced chemical changes rather than surface adhesion effects. To confirm that UV irradiation indeed triggered TT dimer formation inside droplets, we performed immunostaining using anti-TT dimer antibody, which verified TT dimerization in dT40 under these exposure conditions (Figure 1I).

Our quantitative analysis revealed that extended UV exposure (from 30 to 45 min) led to a higher occurrence of doublets and clustered configurations (Figure 2A, and B), as well as a significant increase in droplet-droplet contact area, measured by the chord length (L) (Figure 2A, and C). This reflects slower relaxation due to the expected increase in viscoelasticity and allows doublets with larger chord lengths to be more readily captured during imaging. Additionally, our particle analysis revealed a significantly lower circularity index of +UV45 droplets compared to +UV30 and control (Figure 2D). Size distribution analysis of singlet droplets revealed that the control samples exhibited an exponential distribution (and not log-normal^51^ or power law^52^ behaviours), consistent with a quench-then-coalesce mechanism^53,54^. However, +UV30 samples, and more prominently +UV45 samples, exhibited a clear shift to a power-law distribution, suggesting a transition to an addition of mass growth model^52^ (Figures 2E and S3).

**Figure 2.**
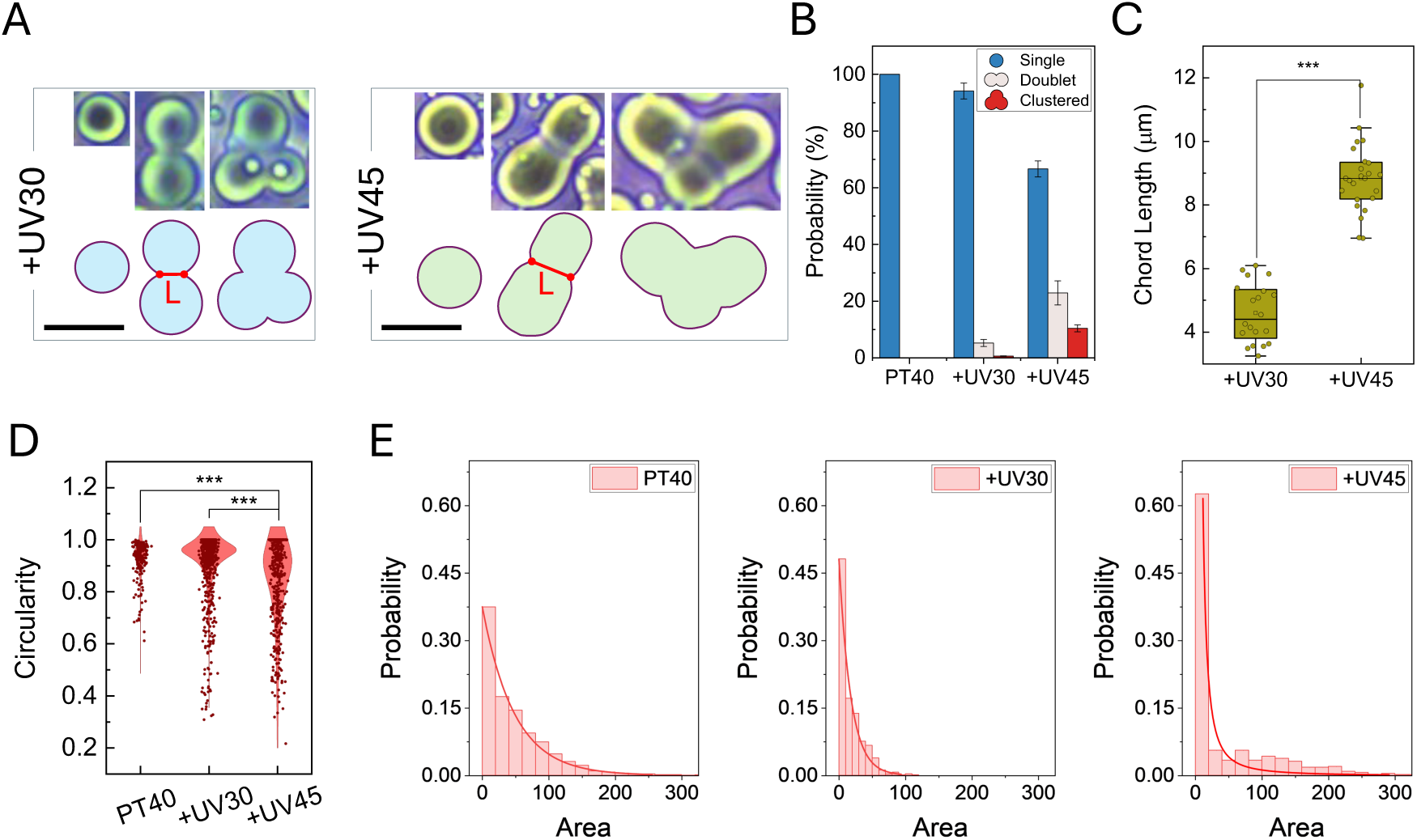
Morphological characterization of PLL:dT40 condensates with and without UV illumination. A) Representative images of single, doublet, and clustered droplets. B) Comparison of droplet morphologies after UV treatments. The longer UV exposure results in increased probability of doublet and clustered configurations formation. Bar plot shows the distribution of droplet shapes; shape percentages were calculated and averaged across three representative fields of view, with error bars indicating standard deviation. C) The longer UV exposure results in increased chord length (L) in doublets. D) Morphological analysis of the droplets by calculating the circularity parameter. Values range from 0 (irregular) to 1 (perfectly circular). E) Size distribution analysis of singlet droplets. Data are fitted by an exponential (*a* + *be^cx^*) and power-law (*ax^b^*) curves. The scale bar represents 10 µm.

### Rheological analysis reveals liquid-to-solid-like transitions upon UV exposure

UV exposure can induce inter- and intra-chain thymine dimers, as confirmed by our spectroscopic and imaging analyses, potentially altering condensate mechanics^55,56^. To investigate this, we performed oscillatory rheology measurements of the storage modulus (G′)and loss modulus (G″) on control and UV-illuminated samples using a recently developed SPM-based approach^22^ (Figure 3A, Note S1). Across the entire frequency range examined (1 Hz to 100 Hz), G″ consistently exceeded G′, indicating that the control droplets exhibited predominantly viscous (liquid-like) characteristics (Figure 3B). We note that the mechanical properties of this condensate system are quite robust within the timescale of our study and do not change over tens of minutes.

**Figure 3.**
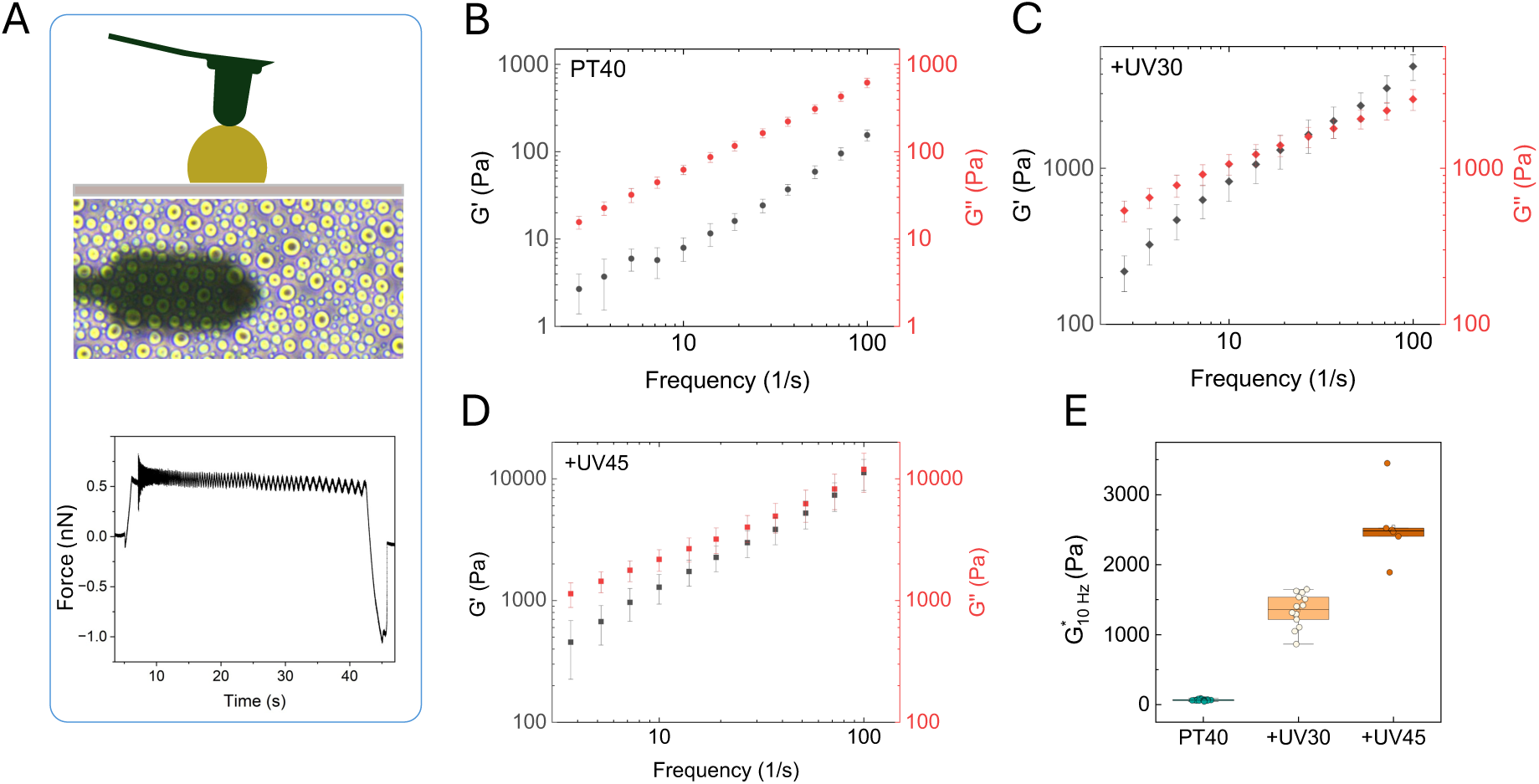
SPM-based mechanical characterization of PLL:dT40 condensates under different conditions (control, +30 min treatment, +40 min treatment). A) Schematic of the experimental setup illustrating the scanning probe contacting a droplet deposited on a substrate. B) Frequency sweep of the storage modulus (G′, black) and loss modulus (G″, red) for the untreated (control) condensate, showing G″ > G′ across the tested frequency range (1 Hz to 100 Hz). C, D) Frequency sweep of G′ (black) and G″ (red) after UV irradiation. E) Complex modulus (G*) plotted at 10 Hz for different conditions, demonstrating a progressive increase in viscoelastic response with increasing irradiation time.

Following UV treatment, significant changes in the viscoelastic responses were observed (Figure 3C). In +UV30 samples, G′ and G″ increased substantially compared to control and intersected around 20 Hz, indicating a liquid-to-solid-like crossover. At frequencies below 20 Hz, G′ rose from 5 Pa to ∼300 Pa, and G″ from 20 Pa to ∼600 Pa. Above 20 Hz, G′ rose more steeply, reaching ∼4500 Pa at 100 Hz, while G″ reached ∼2600 Pa, reflecting dominant elasticity at high frequencies. In +UV45 samples (Figure 3D), a similar trend is seen, but the crossover is shifted to higher frequencies (*f* > 40 Hz), where differences between G′ and G″ become nonsignificant, suggesting the formation of heterogeneous stiff domains. These results support a model in which UV exposure drives condensates from a low-stiffness, liquid-like state to a stiff, solid-like state with slow relaxation at moderate exposure (+UV30), and further UV produces a heterogeneous solid-like system where local stiffening increases elasticity but allows faster relaxation in less-crosslinked regions (+UV45). To better highlight differences between conditions, we also plotted the complex modulus (G*) at 10 Hz (Figure 3E), which reveals a systematic increase in viscoelastic stiffness with increasing irradiation duration. Finally, we measured the terminal viscosity of these condensates at T = 25°C by a linear fit of G″ over frequency. Control samples had the lowest viscosity (6.2 ± 0.09 Pa.s), followed by +UV30 (12.08 ± 0.11 Pa.s), while +UV45 condensates showed the highest (140.77± 11.7 Pa.s).

### UV irradiation significantly alters nucleation and coalescence processes

UV irradiation induces thymine dimerization in DNA molecules, forming intrachain covalent bonds and, with lower probability, interchain crosslinks^49^. This competition between intra- and inter-chain interactions can significantly influence condensate formation and growth. Conversely, the prevalence of each interaction type may depend on the condensation stage, as condensation itself can modulate photochemical outcomes. To investigate this phenomenon, we conducted UV irradiation for 45 minutes under two conditions: the “Premix” condition, where UV illumination was applied to dT40 after the addition of buffer and before the addition of PLL, and the “0 hours” condition, where UV exposure was performed immediately after mixing dT40 and PLL. Interestingly, under the “0 hours” condition, no condensates were detected by our optical microscopic analysis after 4 hours. Instead, a large network of interacting components was formed (Figure 4A, middle panel). Real-time phase-contrast imaging however revealed the transient formation of small condensate-like droplets, which subsequently transitioned into aggregate-like structures during the UV illumination period (Figure S4A). These structures contained DNA and PLL, as shown by fluorescence analysis (Figure S4B). Micro-Flow Imaging (MFI) analysis revealed the real-time formation of small particles with a size distribution distinct from that of pure condensate systems, suggesting the formation of microaggregates with non-circular geometries (Figure S4).

**Figure 4.**
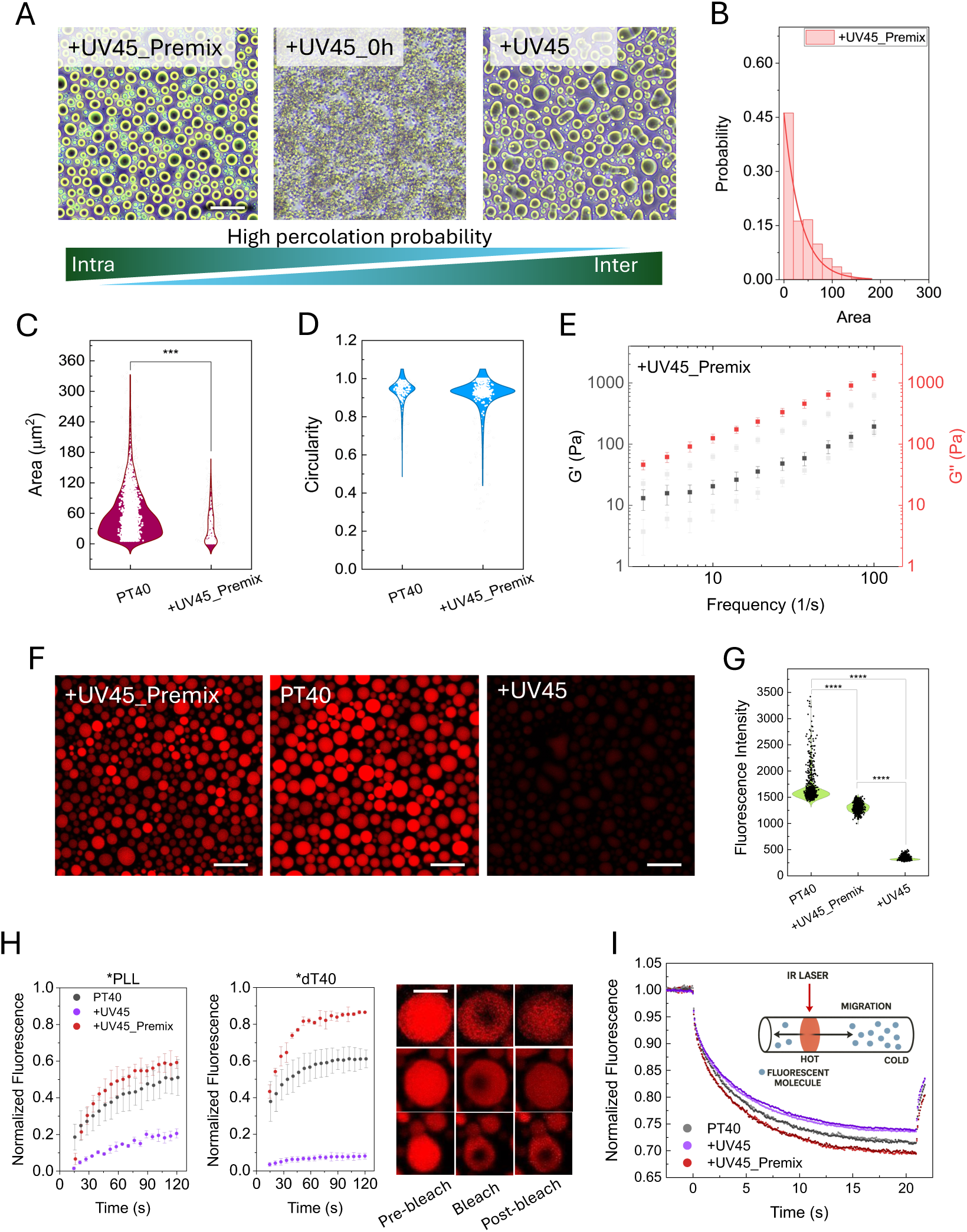
Condensation dramatically impacts the outcome of the UV-induced photochemical reaction in dT40/PLL solution. A Representative phase contrast images show the results of the three different UV exposure protocols: premixing the DNA with buffer before adding PLL (+UV45_Premix), applying UV immediately after mixing (+UV45_0h), and irradiating at 4 h (+UV45_4h). Control samples with no UV illumination are represented as PT40. Early UV illumination (A, +UV45_0h) yields a percolated network rather than distinct droplets, whereas later irradiation (+UV45_4h) produces elongated but well-defined condensates. B) Size distribution analysis of droplets in +UV45_premix samples. Data are fitted by an exponential curve. C, D) Violin plots illustrate droplet morphology and size analysis. E) A frequency sweep of the storage modulus (G′, red) and loss modulus (G″, black). This analysis showed no significant differences between control and premix samples. F) Confocal fluorescence images from the DNA-exchange assay. After droplet formation, 1% TYE-665–labeled dT40 was added to the dilute phase and imaged after 1 h. +UV45_4h shows minimal fluorescence, indicating limited DNA exchange into the crosslinked network, whereas +UV45_Premix and the PT40 control exhibit strong DNA uptake. G) fluorescence intensity analysis of the droplets after addition of the fluorescently tagged dT40 DNA. +UV45_4h samples showed the lowest intensity, indicating minimum penetration of tagged DNA into the condensate. H) Dual-channel FRAP analysis of condensates containing FITC-labeled PLL and TYE-665–labeled dT40 under the three conditions: PT40 (no UV), +UV45_Premix, and +UV45. Premix and control droplets exhibited fast fluorescence recovery for both PLL and dT40, whereas +UV45 condensates displayed negligible recovery in either channel over the experimental timescale, indicating severe immobilization of components following UV-induced DNA crosslinking. Representative pre-bleach, bleach, and post-bleach images represented on the right. I) Microscale Thermophoresis (MST) characterization of DNA extracted from condensates. After UV irradiation, +UV45 condensates were treated with 1 M KCl to dissociate the networks and dissolve the droplet. The released macromolecules from the dilute phase were loaded into MST capillaries and fluorescence was monitored at the loading location. Samples from +UV45 showed the strongest reduction in thermophoretic mobility, PT40 displayed intermediate behaviour, and +UV45_Premix maintained the highest mobility, consistent with the change in the frequencies of inter- (reduced mobility) and intra-chain (enhanced mobility) bond formation. The scale bars represent 20 µm for phase-contrast and confocal images and 5 µm for FRAP analysis images.

Interestingly, in the premix condition, we observed condensate formation (Figure 4A, left), with smaller sizes (Figure 4B, and C) and circularity values comparable to control droplets (Figure 4D), indicating no elongation or deformation. Furthermore, premix UV irradiation had minimal impact on condensate mechanics, compared to the control (Figure 4E), which may be attributed to the low occurrence of inter-chain interactions. Consistent with this interpretation, molecular exchange analysis showed strongly reduced DNA exchange in +UV45, and efficient exchange in +UV45_Premix and control (PT40, droplets received no UV illumination) droplets. To demonstrate this, we added 1% TYE-665 tagged dT40 to the dilute phase after condensate formation and imaged the droplets after an additional hour of incubation. As shown in Figure 4F, the +UV45 samples exhibited the lowest fluorescence intensity compared to both the +UV45_Premix and PT40 samples, with control samples displaying the highest intensity (Figure 4G). Consistent with these observations, FRAP analysis of droplets containing fluorescent components showed rapid DNA fluorescence recovery in both +UV45_Premix and PT40 droplets, but no recovery in +UV45 within the timescale of the experiment (Figure 4H). Finally, Microscale Thermophoresis (MST) performed on these condensate samples revealed significantly reduced mobility in +UV45, consistent with the formation of inter-chain bonds (Figure 4I). Together, these findings support a model in which optically regulated interplay between local or long-range intra-chain and inter-chain interactions governs condensate mechanics.

### UV-stabilized droplets exhibit stability and internal compartmentalization

The above observations suggested that UV exposure cross-links DNA molecules in the condensates, thereby altering condensate mechanics and promoting the formation of arrested fusion droplets, which in turn affects chemical compartmentalization. To further assess stability and compartmentalization, we examined the response of +UV45 droplets to extreme changes in the surrounding environment. This was done by replacing the dilute phase with two extreme conditions that, according to our KCl concentration phase diagram, did not support droplet formation. To do this, following UV irradiation, we removed the original dilute phase and replaced it with pure water (0 M KCl) or an aqueous solution of 1 M KCl. In both cases, the condensates remained stable. In the case of 0 M KCl, phase-contrast microscopy and confocal imaging with FITC-labeled PLL (green) and TYE-665-labeled dT40 (red) revealed the formation of dilute regions within the droplets, indicating an electrostatically driven collapse of the dense network^57^ (darker circles inside the droplets in Figures 5A-D). Washing thus revealed partial compartmentalization induced by UV illumination, which indicates that not all regions were uniformly crosslinked or equally stabilized. Similar dilute compartments have been recently observed in multiple condensate systems upon changes in the ionic strength or temperature^57^. Erkamp et al. showed that this phenomenon occurs when the equilibrium density of the dense phase changes faster than molecular relaxation within the droplet ^57^. Specifically, if the droplet contracts faster than the polymer can diffusively redistribute across the droplet, the contraction will open holes of dilute phase within the dense phase. In our system the removal of salt results in tighter association between dT40 and PLL, favoring contraction of the dense phase network, which occurs much faster than the ∼10 minute timescale required for molecular redistribution (Figure 4H).

**Figure 5.**
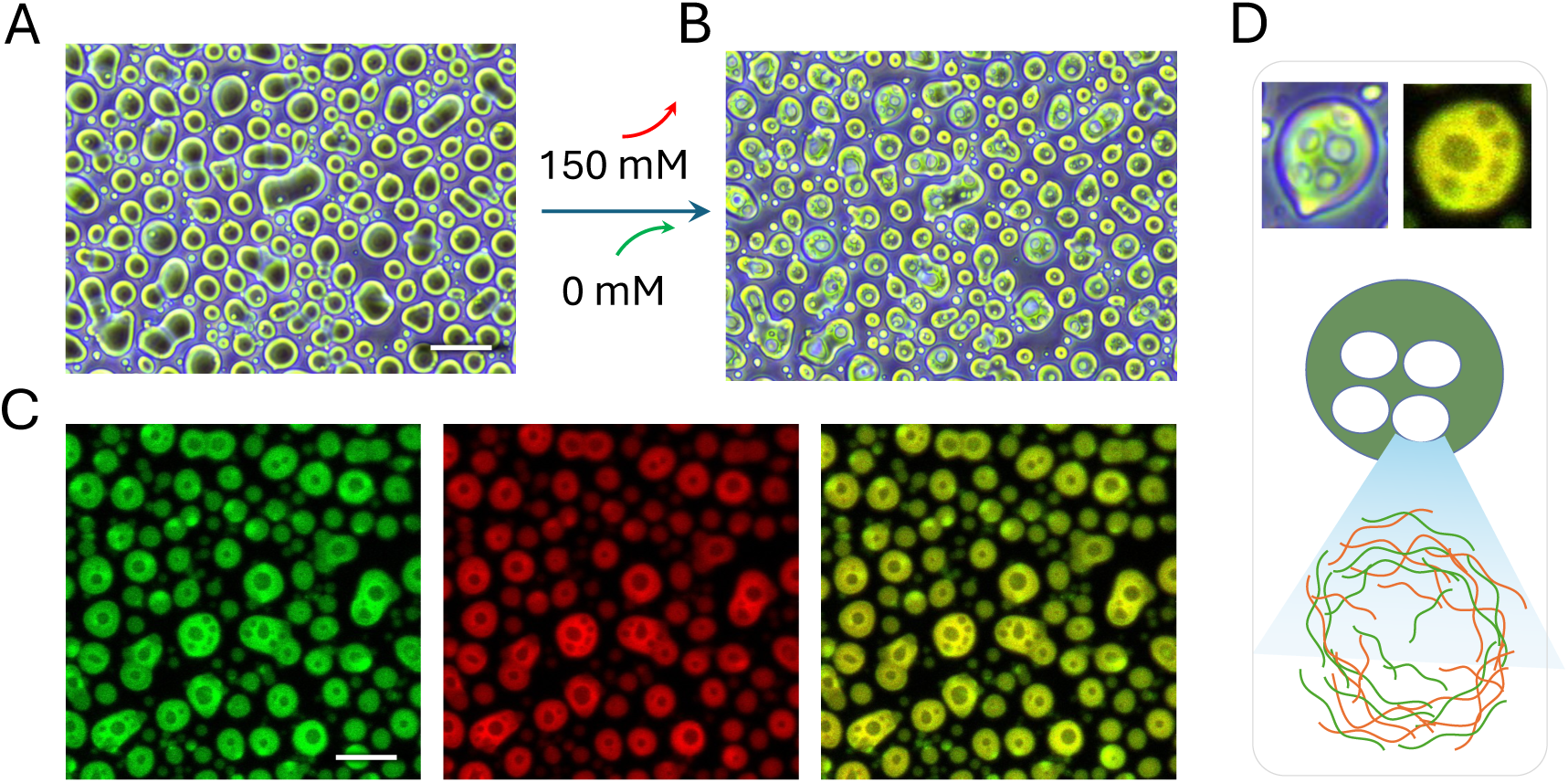
UV irradiated condensate droplets feature internal compartmentalization and remain stable upon drastic changes to the dilute phase. A) Phase-contrast images showing the morphology of +UV45 droplets before and (B) upon exchanging the dilute phase from 150 mM KCl to 0 M KCl. Droplets remained stable but exhibited internal heterogeneity upon this exchange. C) Confocal fluorescence images of +UV45 droplets with labelled FITC-PLL (green) and TYE-DNA (red) in 0 M KCl. D) Magnified view of representative droplets and schematic representation of single droplet, highlighting the observed internal compartmentalization. The scale bars represent 20 µm for phase-contrast and confocal images.

At 1M KCl, confocal fluorescence imaging revealed a marked decrease in overall fluorescence intensity compared to droplets treated with pure water (Figure S5A and B). This reduction indicates a partial release or redistribution of molecular components within the highly screened electrostatic environment. Additionally, unlike the UV-untreated PT40 droplets (Figure S6), fluorescence intensity line scans across individual droplets revealed an interface with high fluorescence intensity (Figure S5 C) in both FITC and TYE channels.

### Direct analysis of droplet fusion dynamics using SPM

Our morphological observations indicated that UV-induced thymine dimerization markedly affects droplet fusion dynamics. To directly quantify these effects, we developed an SPM-based assay for condensate droplet fusion and applied it to control and +UV45 samples. Figure 6A presents the experimental setup for this measurement. A dual-coated Petri dish with BSA and Pluronic F127 (PF127), as described in detail in the methodology section, was used for making the condensate system. Droplets on the Pluronic-coated surfaces were less sticky and more spherical compared to the BSA-coated side. This approach allowed us to easily capture a droplet, move to the BSA side to approach the target droplet, and perform a droplet-droplet fusion experiment (Figure 6B).

**Figure 6.**
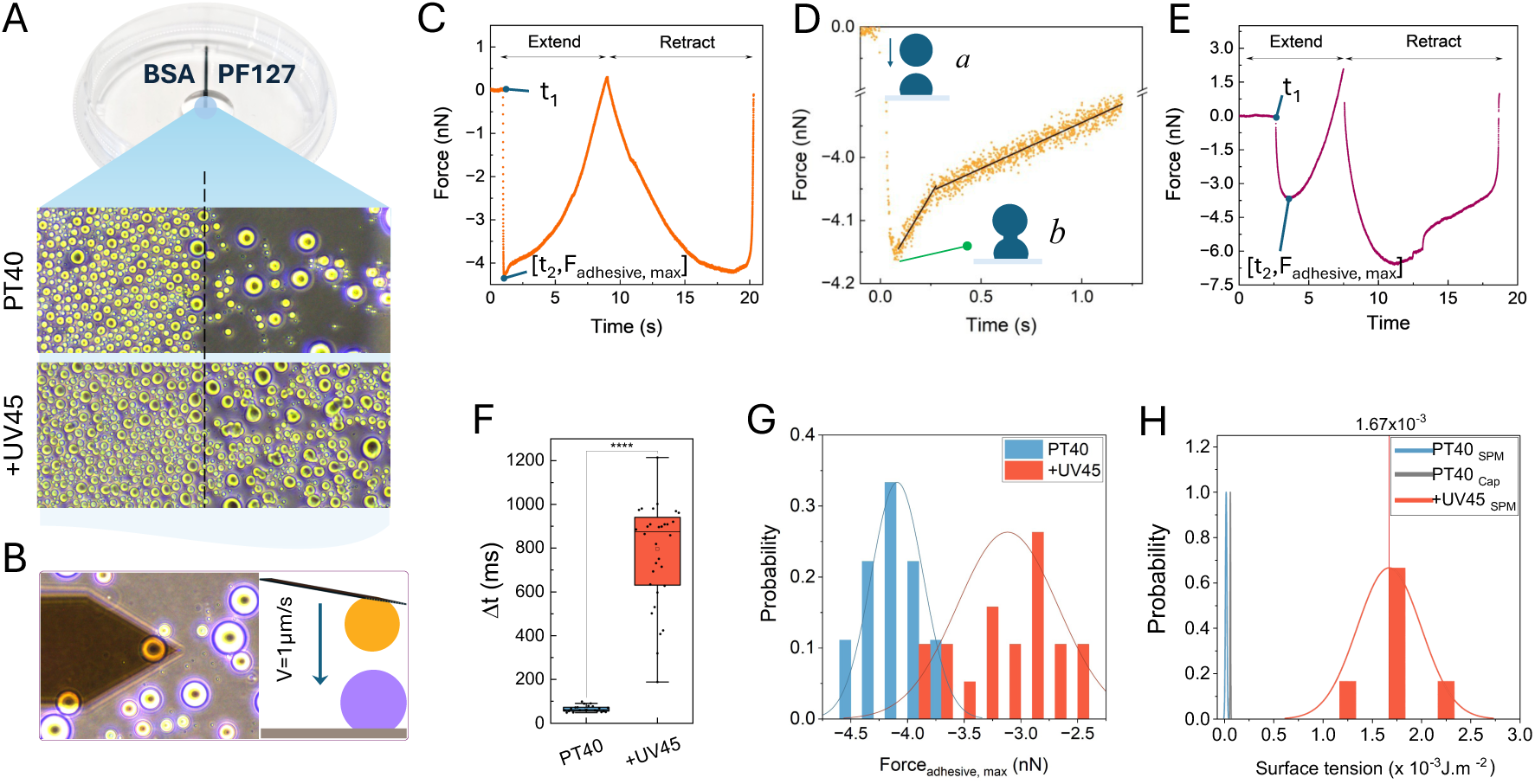
SPM-based droplet–droplet interaction analysis reveals UV-induced changes in fusion dynamics. A) Dual-area coated dish for condensate preparation. Comparison of droplet behaviour on a dual-coated Petri dish with BSA (left) and Pluronic PF127 (PF127; right) surfaces. The black line divides the two coated regions. The lower panels illustrate the differences in droplet formation: droplets on the PA side are more rounded and exhibit reduced stickiness compared to the BSA side, where droplets adhere more strongly to the surface. B) experimental setup for droplet-droplet interaction analysis using SPM. C) Representative example of the recorded Force-time curve for liquid-like PT40 droplets. A cycle includes an extend phase and a retraction phase. During the extend, the piezoelectric actuator is extended, moving the cantilever tip toward the sample surface. The cantilever, with an attached droplet, was moved downward (Extend phase) at 1 µm s⁻¹ until it contacted the droplet on the substrate, consistent with a liquid bridge formation. Following initial contact, the cantilever continued to move downward as droplet fusion progressed to completion, after which it retracted to its original position. The negative portion of the force curve indicates capillary attraction (adhesion) and shows dependency on approach velocity (Figure S7). D) Zoomed in force–time trace showing the interactions between two liquid-like PT40 droplets. The force remains near zero before contact (a), rapidly drops to a negative minimum, suggesting bridge formation (b), and then transitions through two distinct regimes of coalescence and droplet reshaping. E) Representative Force-time trace for +UV45 sample. (F) Box plot comparing the bridge formation times (Δt = t_2_-t_1_) for PT40 and +UV45 samples. The +UV45 sample shows a significantly larger Δt. G) Maximum adhesive force (*F*_adhesive, max_) during droplet interaction for PT40 and +UV45. The +UV45 sample shows a shift toward lower adhesive forces, suggesting that photo crosslinking alters the capillary interactions between droplets by increasing structural resistance to deformation. H) Surface tension (γ) extracted from force–time traces for PT40 based on the capillary model, and from SPM analysis for PT40 and +UV45 samples.

Our workflow began by selecting a target droplet on the PF127 side and imaging it for diameter determination. The cantilever then approached the droplet under a set force of 0.2 nN (waiting time of 0 seconds), which was sufficient for attachment, thereby capturing the droplet. It was subsequently transferred to the BSA side and positioned adjacent to a selected surface droplet. After imaging, the cantilever’s position was adjusted to align the centers of both droplets, and the cantilever was then advanced toward the surface at a constant (“extend”) velocity of 1 µm s⁻¹ (Figure 6B), while force–time traces were continuously recorded.

Figure 6C shows a typical force–time measurement recorded during the interaction of two liquid droplets. Initially, the force remained near zero, indicating that the droplets were not yet in contact. As the cantilever moved downward, the force rapidly shifted (starting at t_1_) to a negative (adhesive) regime when the droplets first came into contact (t_2_), consistent with the formation of a liquid bridge, although direct imaging of the bridge was not possible with our SPM setup. We refer to the time interval between these two events as the liquid bridge formation time (Δt = t_2_-t_1_). After this initial phase, the cantilever continued to move downward at the same speed as the fusion process proceeded to completion. The (liquid) droplets then merged into a single droplet on the substrate. During the subsequent retract phase, the cantilever moved away from the surface, and the force profile remained in the adhesive regime until the connection between the cantilever and merged droplets ultimately broke. Once detached, the cantilever returned “dry” to its original position, leaving an enlarged droplet on the substrate.

Notably, our data revealed a significant difference in the force-time profiles of UV-treated and untreated samples. In non-crosslinked liquid-like droplets, the force drops instantly and then exhibits a two-stage recovery (Figure 6D) after reaching the negative minimum: an initial rapid increase due to fast bridge expansion, followed by a slower phase associated with droplet relaxation and substrate wetting. We observed significantly shorter Δt (Figure 6F) and a higher maximum adhesive force (Figure 6G) in the fusion of control droplets. By contrast, in +UV45 droplets with gel-like behaviour, the force decreases gradually, and the distinct two-regime behaviour is absent; instead, the force recovery occurs gradually and in a continuous manner (Figures 6E, S8).

To examine the geometric scaling expected for capillary coalescence, we compared *F*_adhesive,max_ with the reduced radius *R*^∗^ = *R*_1_*R*_2_/(*R*_1_ + *R*_2_) for each droplet pair ^58^. Control droplets followed the linear relation *F*_adhesive,max_ = 4*πγR*^∗^, yielding an apparent interfacial tension (Fig. 6H and S9). UV-treated droplets systematically deviated below this line, which can be attributed to the viscoelastic contributions, not included in the simple capillary model of pure liquids. The observed deviation from the capillary limit can be rationalized by considering that crosslinking introduces an additional mechanical resistance that reduces effective adhesion. To represent these viscoelastic contributions in a compact phenomenological way, we introduced an empirical modulation factor *S*(*G*’, *G*”), yielding

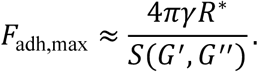

For un-crosslinked capillary liquids, *S* ≈ 1, whereas in viscoelastic droplets *S* > 1 increases with storage and loss moduli. Several functional forms for *S* reproduce the measured trend, including a rational form

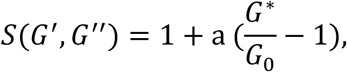

(with 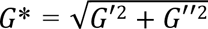 and smooth sigmoidal crossover variants, including a logistic form (Note S2).. While not intended as full mechanical models, these phenomenological descriptions capture the continuous transition from a capillary-dominated (surface tension driven) to a viscoelastic-dominated regime in an internally consistent manner. *G*_0_ represents the corresponding modulus of the un-crosslinked liquid, providing a dimensionless scaling. The approach is most robust under conditions of slow coalescence, a regime that is characteristic of viscoelastic systems^59^. When applied to other systems, the same framework can be adapted by adjusting parameters to match the characteristic crossover behaviour of different condensate chemistries.

We note that the *γ* values calculated from rheology closely agree with those extracted from the capillary model (Figure 6H) for control droplets; however, the measured values are slightly smaller, which can be attributed to mild facilitation of fusion due to favorable molecular interactions in condensates as compared to pure capillary coalescence (accounted for by *S*_0_ in the model, see Note S2). Overall, our observations suggest a model in which the gel-like+UV45 droplets exhibit fusion behaviour that is strongly influenced by viscoelasticity (controlling the absolute fusion force), while the heterogeneity in the normalized fusion force reflects contributions from fluctuations in surface tension as well (See Note S3).

Finally, we examined whether the balance between the occurrence of inter- and intra-chain TT dimers affects fusion dynamics. Our rheology analysis showed that in the regime dominated by intra-chain contacts, as seen in +UV45_Premix, both storage and loss moduli are substantially smaller than when inter-chain contacts are frequent, as in the case of +UV45 (Figure S10). Furthermore, surface tension follows a similar trend, with +UV45 droplets exhibiting higher surface tension. Given that 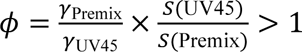, the absolute value of the adhesive force is expected to be higher in +UV45_Premix. Consistent with this, our experimental analysis showed that adhesive force is reduced for +UV45, and the slow structural relaxation of its crosslinked network leads to a marked increase in Δt (Figure S10, and 7). Moreover, the decrease in Ψ from Premix to +UV45 (observed in both models; See Note S3) is at least 25%, suggesting that accumulation of inter-chain crosslinks suppresses the sensitivity of fusion forces to viscoelastic fluctuations (consistent with the effective saturation of the moduli), thereby shifting the dominant source of variability toward interfacial mechanics.

## Discussion

Mechanics of phase-separated biomolecular condensate droplets can be altered in various ways, including by changing their chemical content, internal structure, or environmental temperature. Here, we showed how light can be leveraged to program the connectivity and internal organization of a nucleic acid-based condensate system, thereby changing condensate mechanics with no alterations to atomic (elemental) composition. UV exposure drives a controlled transition from liquid-like to gel-like droplet states, altering condensate morphology, structural mechanics, and fusion kinetics. This is driven by the formation of intra- and inter-chain bonds that contribute to a dramatic increase in viscous and elastic moduli, stiffen the droplets towards gel-like/solid-like states, and shift the fusion process from an interfacial tension-driven fast process to a viscoelastic-dominated slow fusion regime, leading to the observation of arrested fusion droplets. Despite transition towards solid-like states under extreme crosslinking conditions, the system may relax on short time scales, as the organization of the dense domain and surrounding dilute phase enables fast stress relaxation.

Our study presents a concrete example of a photochemical process being modulated by condensation, as the balance between the frequencies of inter- and intra-chain interactions depends on the presence and stages of condensation (Figure 7). Irradiating the system immediately after mixing the building blocks instead yields a network-like structure^60^. In contrast, applying UV after droplets had formed led to rearrangements primarily within the droplet interior, resulting in condensate mechanical transitions. The premix condition, where the nucleic acid was irradiated prior to adding PLL, produced smaller droplets that retained a largely viscous profile, presumably because intra-chain bond formation dominated in the absence of the polypeptide partner, thus limiting the interaction sites for PLL to form larger complexes and grow. Our observations can be explained by noting that (1) intra-chain bonds bend the DNA, reduce its radius of gyration, and stiffen the polymer, and (2) condensation affects the radius of gyration of polymers as well as the density and inter-chain contact probability ^16,61–63^, which will in turn regulate the occurrence of short-range and long-range intra-chain and inter-chain thymine dimers due to topological effects and steric constraints.

**Figure 7.**
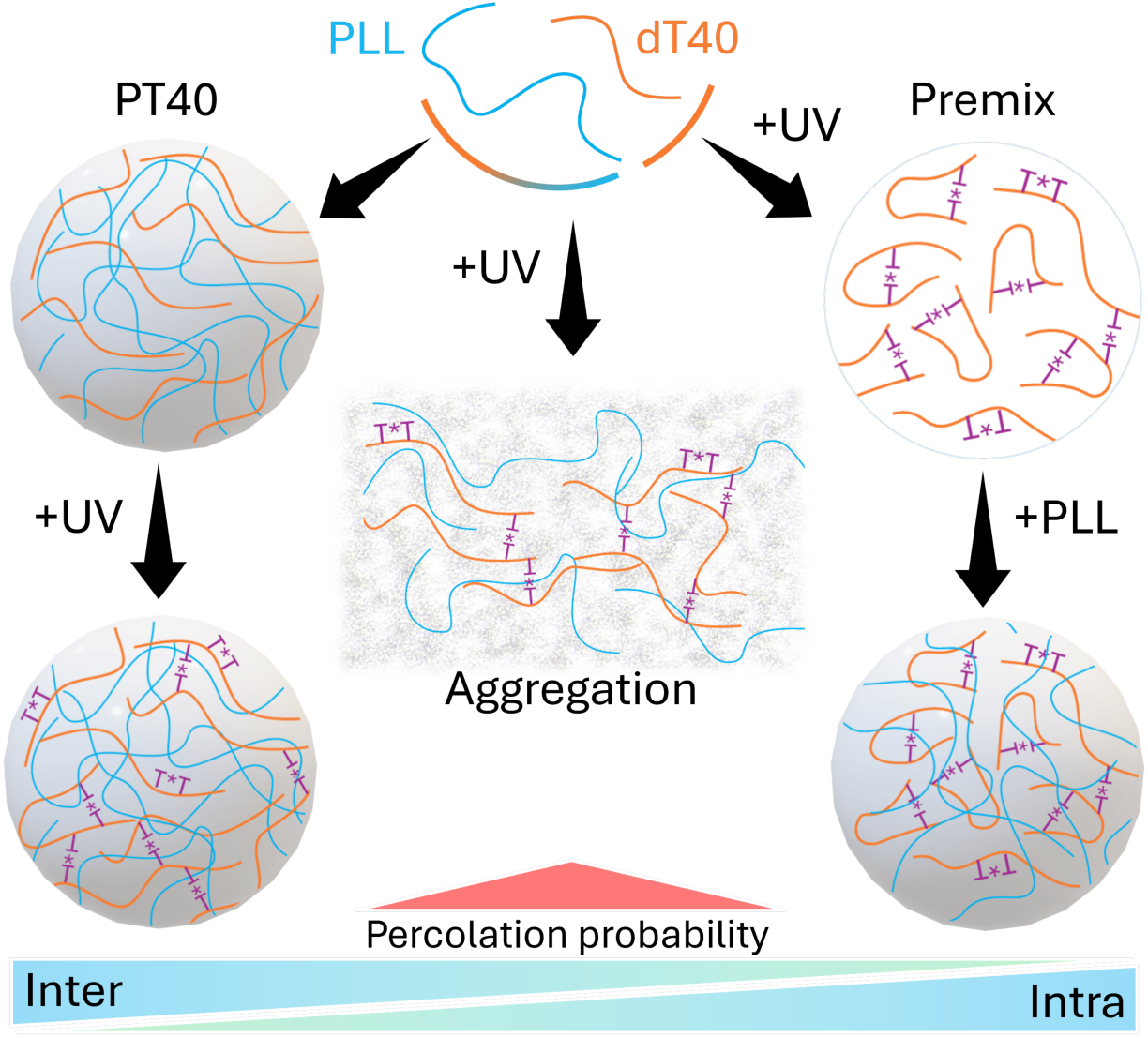
Reciprocal interplay between photochemical reactions and condensation, governed by shifts in inter- and intra-chain interactions. Our data indicate the dominant role of the intra-molecular interactions in premix conditions, which decreases in samples exposed to UV after condensates formed, mainly due to the presence of interacting partners in the environment and increases in dT40 gyration radius that dominated inter-molecular interactions. Between the two extremes, UV irradiation leads to the formation of a percolated network of interacting polymers in 0-hour condition, which results in the formation of aggregates with globally connected chains. The red triangle denotes an increased probability of network percolation in 0-hour UV-illuminated samples.

Our finding provides insights into the role that UV illumination may have played in the origin of life. Early Earth conditions likely exposed primitive protocells to variable doses of solar UV radiation, which could have simultaneously driven molecular crosslinking (through pyrimidine dimerization in early nucleic acids^64,65^) while also posing a threat to nucleic acid stability^66^. Recent models suggest that compartmentalization may have provided a protective niche that shielded nascent genetic material from harmful UV doses^67^. Notably, our findings reveal that UV illumination induces compartmentalization and enhances the stability of the coacervate droplets against drastic changes in environmental chemistry. This mechanism may have played a key role in the resilience and evolution of early biopolymeric systems, balancing the harmful and beneficial effects of radiation on the development of life-like systems.

This study provides new tools and approaches for the synthesis and characterization of materials. The SPM-based assay presented here enables the analysis of condensate droplet fusion dynamics under physiological and pathological conditions. SPM offers straightforward force calibration, and its open geometry makes it possible to bring separately treated droplets into controlled contact and then characterize the merged object, capabilities that are difficult to achieve with other methods. The proposed suppression model, despite its simplicity, captures the interplay between surface tension and viscoelasticity reasonably well and can be applied to a wide range of crosslinkable condensate systems, although more elaborate phenomenological forms (e.g., multi-parameter crossover) and models with explicit microscopic grounding could be considered in more complex scenarios. Furthermore, the synthesis of stabilized condensate droplets with tunable mechanical properties may open new avenues in materials and particle synthesis for biomedical and engineering applications. Such tunability offers exciting opportunities for the design of programmable biomolecular materials, especially given the contact-free nature and experimental convenience of UV-induced crosslinking^68^. The approach presented in this study is versatile and can be readily applied to other polyelectrolyte–nucleic acid systems or under different environmental conditions. It thus opens avenues for engineering synthetic organelles and soft materials that dynamically respond to changes in their environment.

## Acknowledgments

The authors thank Sander Woutersen (University of Amsterdam) for valuable discussions. The authors also gratefully acknowledge the support of the Leiden Cell Observatory (LCO). In particular, we thank Kostas Tassis of the LCO for his assistance with this work.

## Author Contributions

Conceptualization, A.M. and V.S.; methodology, A.M. and V.S.; investigation, V.S., A.M., D.B., J.S.; visualization, V.S.; formal analysis, V.S. and F.W.; writing original draft, V.S. and A.M.; writing review & editing, V.S., A.M., J.S., D.B., F.W.; project administration, funding acquisition, and supervision, A.M.

## Data Availability

The data that support the findings of this study are available from the corresponding author upon reasonable request.

