## Supplementary Information for "Optically driven control of mechanochemistry and fusion dynamics of biomolecular condensates via thymine dimerization"

### Supplementary Figures

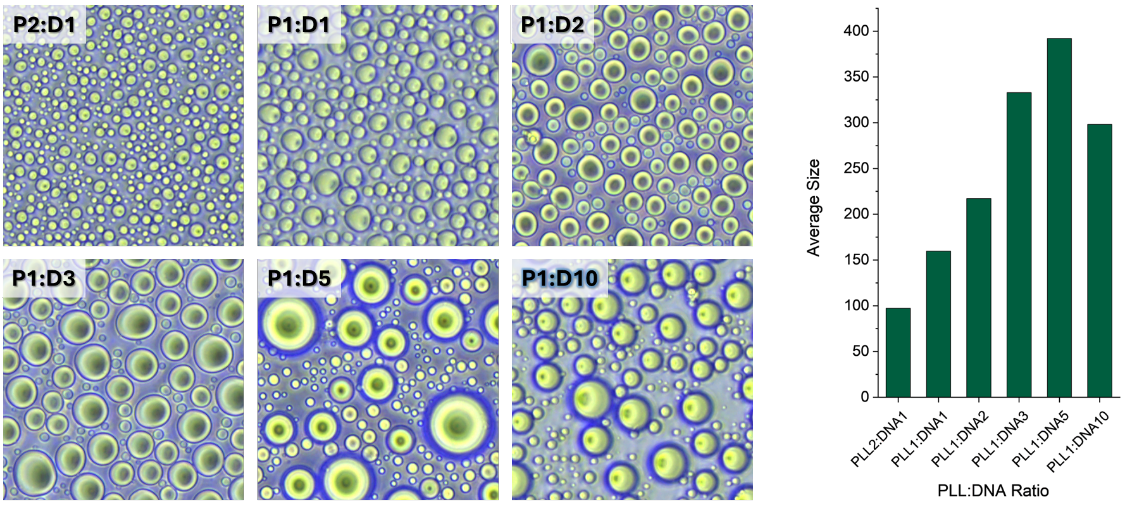


*Figure S 1. Microscopy images showing condensate formation at different DNA-to-PLL ratios. Droplet size increases as the DNA concentration increases relative to PLL. The 1:1 (P1:D1) ratio was selected as the optimal condition due to its suitable average droplet size for mechanical analyses, including rheological measurements and droplet-droplet interaction studies. The bar graph on the right quantifies the trend of droplet size variation across different conditions.*

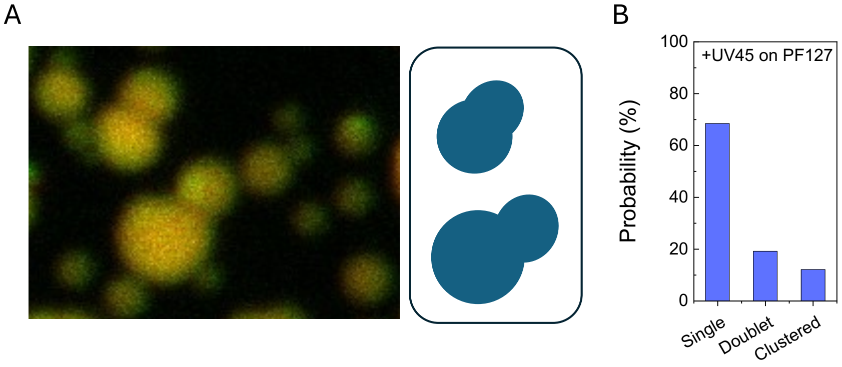


*Figure S 2. A) Emergence of fusion arrest state of the droplets after 45 minutes of UV irradiation on a PF127 coated surface. FITC-labeled PLL is shown in green, TYE-665–labeled dT40 is shown in red, and their overlap appears yellow. B) Probability of doublet and clustered configurations formation showed a good agreement with the formation of the same configurations on the BSA passivated surface.*

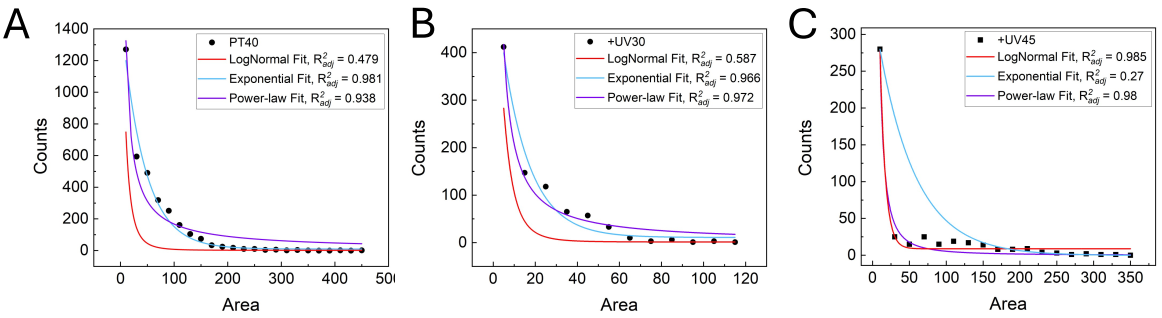


*Figure S 3. Size distribution analysis of droplets fitted with Log-Normal, Exponential, and Power-Law models. The control (PT40) sample followed an exponential pattern, whereas the UV-treated samples tended to follow power-law distributions. The +UV30 sample fit both the exponential and power-law models reasonably well, while +UV45 followed the power-law model. Theoretical arguments for systems governed by fragmentation and coalescence generally point toward log-normal size distributions. This follows from the multiplicative nature of random breakup or coalescence steps: when structures repeatedly divide or merge in a stochastic, multiplicative way, the central limit theorem tends to produce a log-normal distribution. In that sense, observing a log-normal distribution would indicate that condensate sizes are set mainly by random multiplicative breakup, whereas deviations suggest that other mechanisms dominate. In our system, none of the measured distributions were log-normal, indicating that simple multiplicative breakup does not control droplet sizes. Before UV illumination, we consistently observe an exponential distribution, which is consistent with fast nucleation followed by slower coalescence: droplets appear rapidly and then grow primarily by merging, naturally giving rise to an exponential form. Under UV illumination, increased crosslinking alters the growth pathway. Coalescence becomes increasingly arrested while the structures themselves become nonspherical and develop surface areas larger than spheres of equal volume. Their enlarged, irregular surfaces increase their likelihood of additional encounters, effectively creating preferential attachment. This mechanism can explain the transition to a power-law (heavy-tailed, scale-free) size distribution after long illumination times. At intermediate UV exposure, the system shows features of both regimes, reflecting a gradual shift from simple coalescence dynamics to growth dominated by UV-induced preferential attachment.*

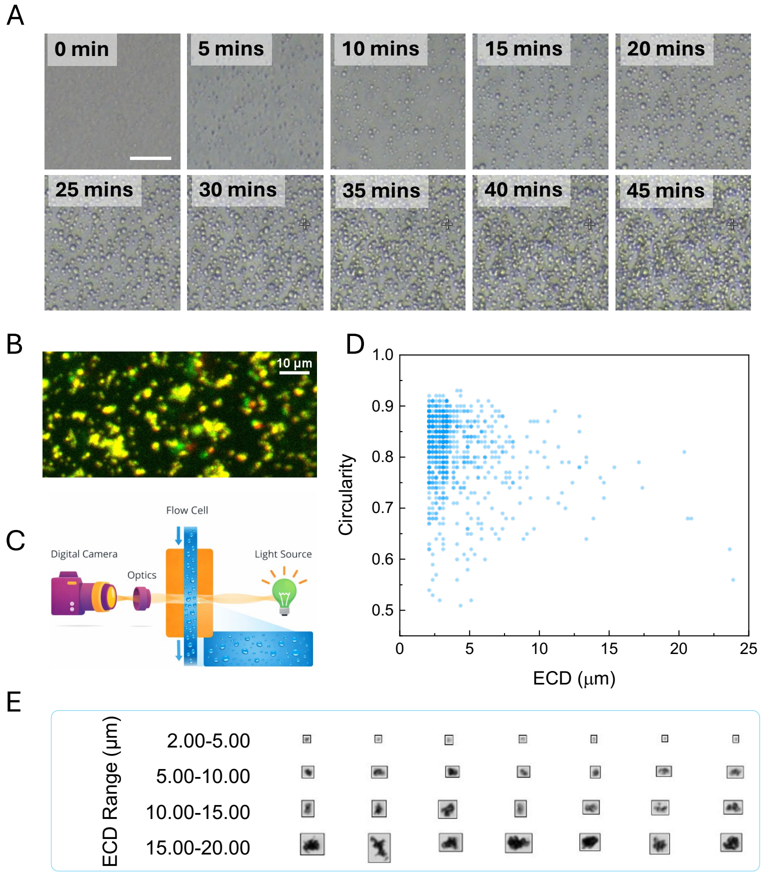


*Figure S 4. Real-time observation of +UV45_0h aggregate-state formation. A) Real-time phase-contrast microscopy images acquired during UV irradiation from 0 to 45 minutes after mixing the two components. Small condensate-like droplets appeared and progressively formed a network-like structure. The analysis suggests condensate-to-aggregate transition as a contributing mechanism to observed behaviour of +UV45_0h state. B) Fluorescence microscopy image of the UV45_0h sample captured immediately after 45 minutes of UV irradiation, prior to mixing the two components, illustrating formation of aggregate-like structures. FITC-labeled PLL is shown in green, TYE-665–labeled dT40 is shown in red, and their overlap appears yellow. C–E) Micro-Flow Imaging (MFI) of particles formed in +UV45_0h samples. Samples were collected at t = 45 min; particle distributions reflect the state at sampling and during MFI acquisition. C) Schematic of the Micro-Flow Imaging (MFI) setup. In this technique, the sample flows through a narrow imaging cell while a bright-field light source illuminates the particles. As particles pass through the detection zone, a high-resolution digital camera captures individual images in real time. MFI identifies and quantifies particles based on their silhouette, enabling simultaneous measurement of size, shape, and morphological features without relying on scattering or fluorescence. D) Scatter plot of particle circularity versus equivalent circular diameter (ECD), revealing a broad distribution of morphologies characteristic of early-stage aggregation. E) Representative MFI particle images binned into four ECD ranges, illustrating the transition from smaller, more circular species to larger and increasingly irregular aggregates. Scale bar represents 10 µm.*

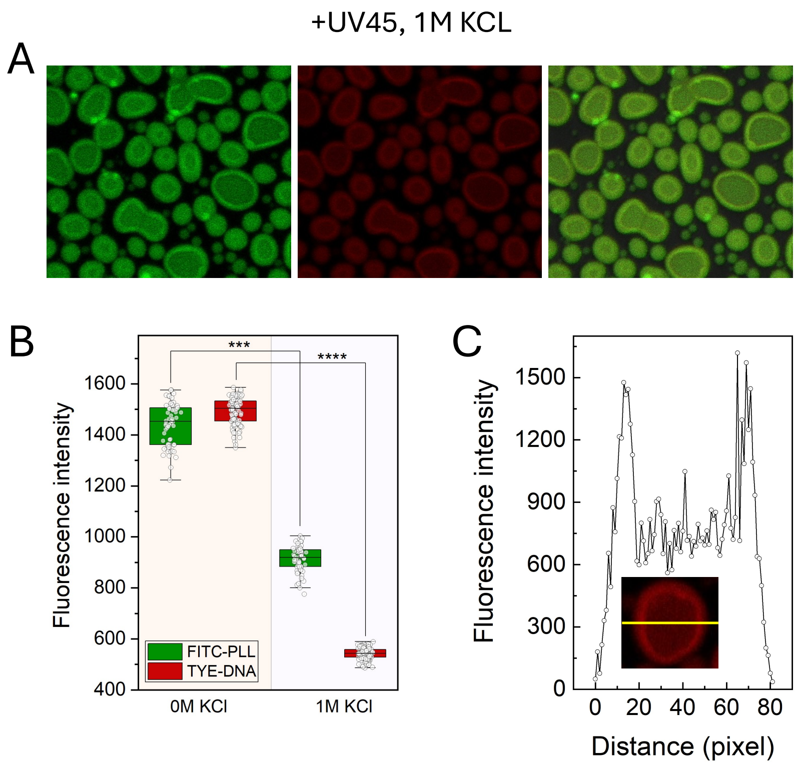


*Figure S 5. A) Confocal fluorescence images of +UV45 droplets with labeled FITC-PLL (green) and TYE-DNA (red) in 1 M KCl. The fluorescence intensity distribution suggests differential retention of labeled components, with TYE-DNA fluorescence being notably reduced in high salt conditions. B) Box plot analysis of fluorescence intensities in 0 M KCl, showing comparable retention of FITC-PLL and TYE-DNA, suggesting minimal molecular segregation under these conditions, while in 1 M KCl, revealing a significant reduction in TYE-DNA fluorescence compared to FITC-PLL, indicating preferential retention of PLL and selective loss of DNA molecules in high-salt conditions. C) Line scan fluorescence intensity profiles of droplets in 1 M KCl, showing differences in fluorescence distribution across a droplet. It indicates a higher intensity of the components at the edge of the droplets.*

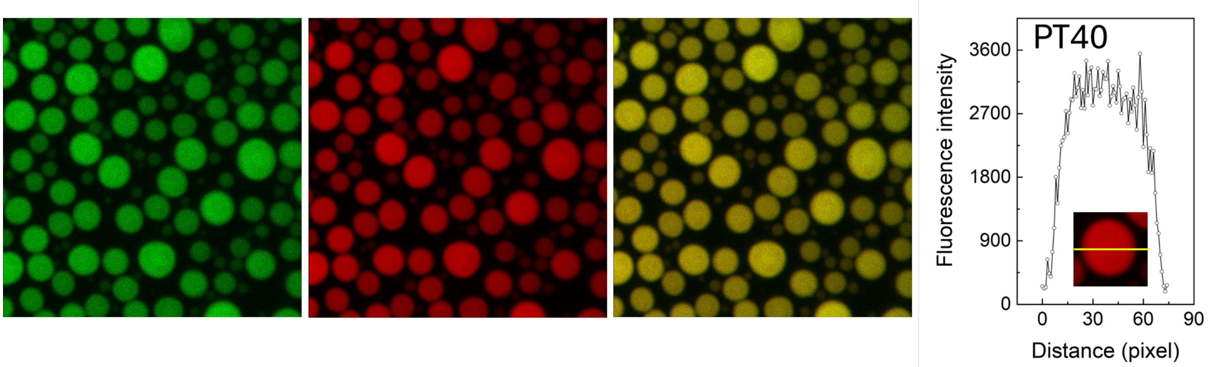


*Figure S 6. Fluorescence microscopy images of PT40 droplets in green and red channels, with the merged image (right) showing uniform fluorescence distribution. The fluorescence intensity profile (far right) confirms the homogeneous distribution of fluorescence across the droplet, indicating no significant compartmentalization in non-UV-treated samples.*

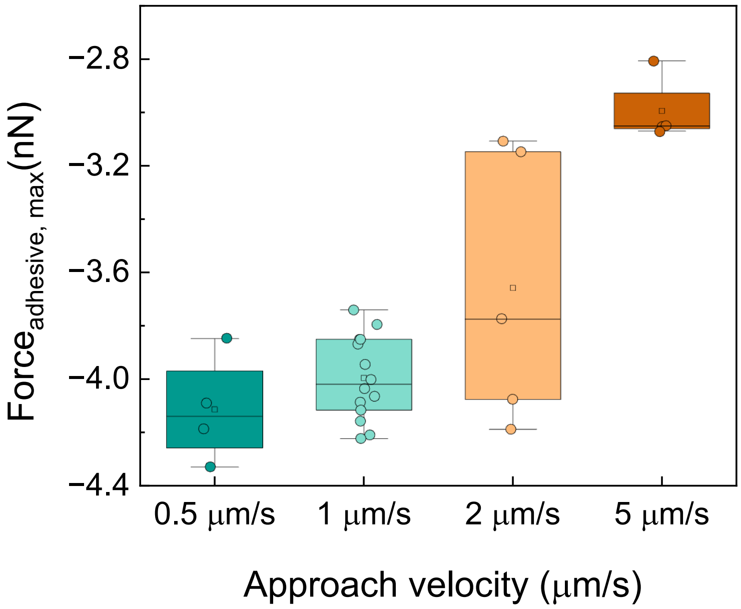


*Figure S 7. Adhesive force decreases with increasing droplet approach velocity. The cantilever approached the surface droplet at four different velocities of 0.5, 1, 2, and 5 µm s$^{-1}$, and the resulting maximum adhesive forces were calculated from the force time traces. Our data showed $F_{adhesive,max}$ decreased systematically from roughly 4.3 nN at 0.5 µm s$^{-1}$ to 3.0 nN at 5 µm s$^{-1}$.*

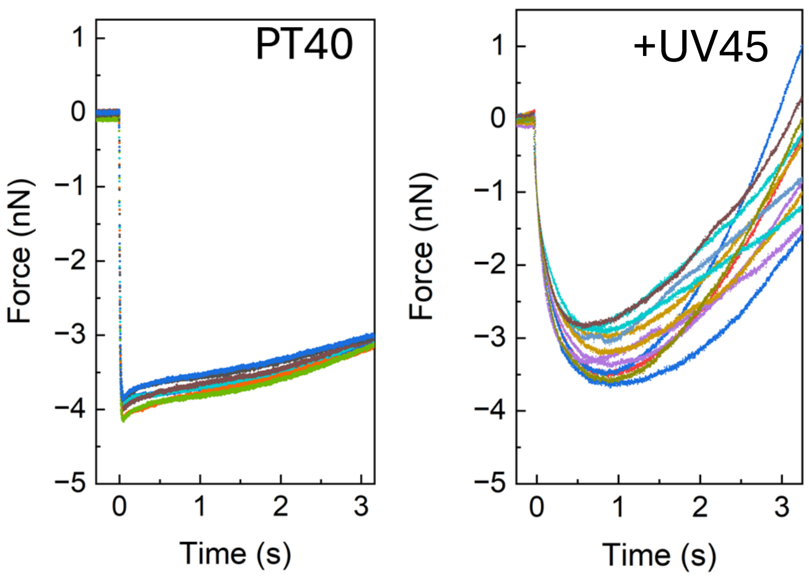


*Figure S 8. Force-time traces from droplet-droplet interaction experiments, demonstrating the reproducibility of force measurements across different droplet pairs. The traces highlight the consistency in interaction dynamics, including the characteristic time and maximum adhesive force for both PT40 and +UV45 samples.*

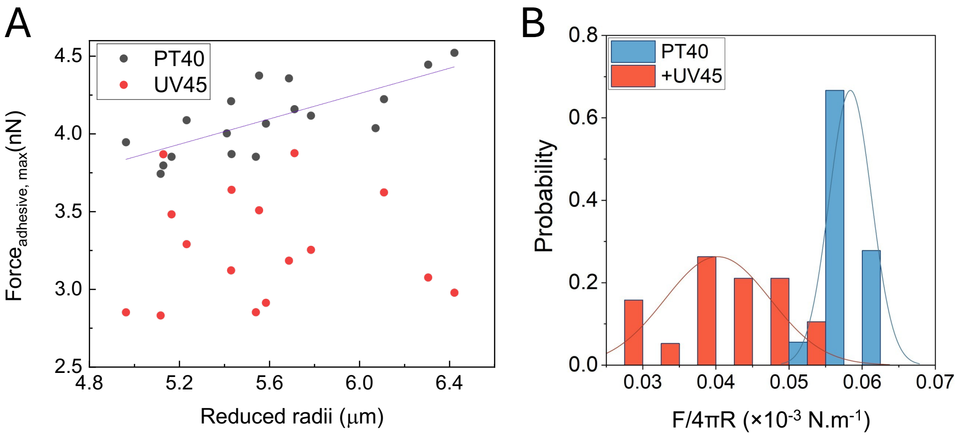


*Figure S 9.* Geometric scaling of adhesive forces and viscoelastic suppression of capillary coalescence. *A) Maximum adhesive force $F_{adh,max}$ plotted against the reduced radius $R = \frac{R_1 R_2}{R_1 + R_2}$ for individual droplet pairs. Control droplets (gray) follow the expected capillary scaling $F_{adhesive,max} = 4\pi\gamma R$, yielding an apparent interfacial tension from the linear fit. UV-treated droplets (red) systematically fall below this capillary limit, indicating reduced effective adhesion due to viscoelastic resistance introduced by UV-mediated crosslinking. B) Probability distributions of the normalized force $F/4\pi R$ for control PT40 and UV-treated (+UV45) droplets. Control droplets cluster near a single $\gamma$ value, consistent with a simple liquid interface, whereas UV-treated droplets show broadening and a shift toward smaller values, reflecting viscoelastic suppression of adhesion.*

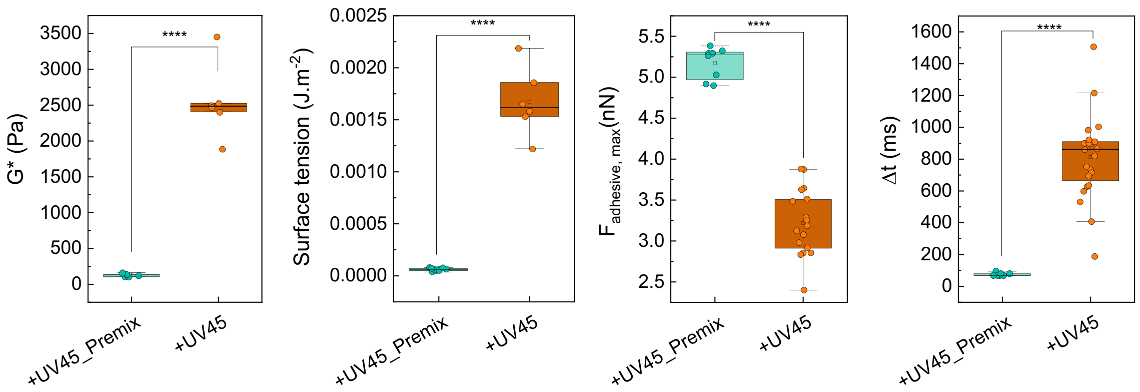


*Figure S 10. Comparative analysis of droplet mechanics and fusion dynamics between the +UV45_Premix condition, where intra-chain formation dominates, and inter-chain events are rare, and the +UV45 droplets, where inter- and intra-chain formations occur frequently.*

### Supplementary Tables

**Table S1. Experimental inputs and parameter values used for viscoelastic modulation (VEM) models.** To apply the VEM models, we used condition-averaged $\gamma$ and $G^*$ (evaluated at 10 Hz, corresponding to the deformation rate from the 1 μm $s^{-1}$ extend velocity) obtained from our rheology study to estimate model parameters.

| Model inputs and parameters | Value (Mean ± STDEV) | Value (Median) | Condition |
|---|---|---|---|
| $G'$ | 1286.24 ± 342.08 | 1230.39 | +UV45, at 10 Hz |
| $G''$ | 2177.52 ± 459.97 | 2167.28 | +UV45, at 10 Hz |
| $G'$ | 8.53 ± 3.51 | 8.22 | PT40, at 10 Hz |
| $G''$ | 63.38 ± 4.63 | 61.85 | PT40, at 10 Hz |
| $\gamma$ | 0.00167 ± 0.00032 | 0.00166 | +UV45 (SPM) |
| $\gamma$ | 1.607277e-05 ± 7.388583e-06 | 1.362670e-05 | PT40 (SPM) |

### Supplementary Notes

#### Note S1. Rheology analysis of the droplets using SPM

Oscillatory rheology measurements of the storage modulus ($G'$) and loss modulus ($G''$) were performed at the frequency range from 1 Hz to 100 Hz on control and UV-illuminated samples. For doing this, the SPM cantilever was positioned above the droplet, and then mechanical manipulation steps started, including indentation with a constant velocity of 1 um/s until reaching the force set point of 0.3 nN, 1s relaxation time, and then 14 consecutive oscillations. After each manipulation cantilever detached from the droplet and retracted to its initial position. Figure 3A represents the schematic and real microscopic view of the experimental setup. Experiments were performed in triplicate with n=10 number of droplets per replicate. Each replicate was a fresh preparation on a different day. Acquired data were analyzed using a custom-made code, and rheological parameters were calculated.

#### Note S2. Phenomenological Viscoelastic Modulation Models for Droplet Fusion

To represent deviations from the ideal capillary limit in a dimensionless, phenomenological framework, we introduce a viscoelastic modulation factor, $S(G', G'')$, that captures the modulation of the fusion force by droplet properties. The models recapitulate the continuous transition from capillary-dominated to viscoelastic-dominated behavior in condensate systems upon crosslinking.

For simplicity, we consider a linear (rational) form,

$$S(G', G'') = S_0 + \alpha\left(\frac{G^*}{G_0} - 1\right)$$

where $S_0$ represents the minimal modulation in the un-crosslinked system, $G^* = \sqrt{G'^2 + G''^2}$, $G_0 = G_{\mathrm{PT40}} = 63.96$, and $\alpha$ describes the increase of suppression with modulus. Enforcing exact agreement with the measured modulation factors at the experimentally defined modulation for PT40 and +UV45, yields $\alpha = 1.0207$. For a more refined description with a continuous S-shaped crossover, we introduce a logistic model,

$$S(x) = S_0 + \frac{S_\infty - S_0}{1 + \exp\left[-\alpha(x - \beta)\right]}, x = \frac{G^*}{G_0},$$

with the experimentally measured plateaus $S_0 = 0.2725$ and $S_\infty = 39.63$. Using a tolerance-based approach ($\delta = 0.01$), the logistic parameters are $\alpha = 0.2383, \beta = 20.28$, ensuring a smooth, robust crossover. The sigmoid model provides a higher-fidelity description of viscoelastic modulation, compared to the simple linear (rational) model.

The fusion force is then expressed as

$$F_{\mathrm{adh,max}} \approx \frac{4\pi\gamma R^*}{S(G', G'')},$$

where $R^*$is the reduced radius of the droplet pair.

**Note S3. Error Propagation Analysis for Fusion Mechanics**

To quantify the relative contributions of surface tension and viscoelasticity to the variability of the fusion force, we consider the normalized quantity

$$f = \frac{F}{4\pi R^*} = \frac{\gamma}{S(G^*, \alpha)},$$

which isolates the material parameters by removing the trivial geometric dependence on droplet size. The spread of the measured $f$ values follows directly from error propagation,

$$\sigma_f^2 \approx \left(\frac{\partial f}{\partial \gamma}\right)^2 \sigma_\gamma^2 + \left(\frac{\partial f}{\partial G^*}\right)^2 \sigma_{G^*}^2.$$

The relative weights of these terms determine how much of the observed variability in the fusion force originates from fluctuations in interfacial tension versus fluctuations in viscoelasticity. To express this balance in dimensionless form, we define

$$\Psi \equiv \left|\frac{(\partial f/\,\partial G^*)\,\sigma_{G^*}}{(\partial f/\,\partial \gamma)\,\sigma_\gamma}\right|,$$

so that $\Psi \gg 1$ indicates that variability in $G^*$contributes dominantly to the spread of the measured fusion forces, while $\Psi \ll 1$ indicates dominance of surface-tension fluctuations.

Explicitly,

$$\Psi = \frac{\sigma_{G^*}}{\sigma_\gamma} \frac{\gamma}{S} \left(\frac{\partial S}{\partial G^*}\right),$$

with

$$\sigma_{G^*} \approx \sqrt{\left(\frac{G'}{G^*}\sigma_{G'}\right)^2 + \left(\frac{G''}{G^*}\sigma_{G''}\right)^2}\ .$$

For any form of $S$, the resulting expressions for $\Psi$ depend only on measurable quantities $(\gamma, G^*, \sigma_{G^*}, \sigma_\gamma)$ (See Table S1) and on the model parameters $(\alpha, \beta, G_0)$. Across both the linear (rational) and logistic viscoelastic-modulation models, the error-propagation factor remains of order unity for +UV45 system. Consequently, uncertainties in $\gamma$ consistently make a non-negligible contribution to the overall heterogeneity in $f$ for UV-treated samples, regardless of the specific viscoelastic modulation model used. Viscoelasticity dictates the absolute scale of $f$, but variability in $\gamma$ remains comparable to, or larger than, the propagated variability from $G^*$.
